# The impact of interactions on invasion and colonization resistance in microbial communities

**DOI:** 10.1101/2020.06.11.146571

**Authors:** Helen Kurkjian, M. Javad Akbari, Babak Momeni

**Affiliations:** Department of Biology, Boston College, Chestnut Hill, MA 02467

## Abstract

In human microbiota, the prevention or promotion of invasions can be crucial to human health. Invasion outcomes, in turn, are impacted by the composition of resident communities and interactions among resident microbes. Microbial communities differ from communities composed of other types of organisms in that many microbial interactions are mediated by chemicals that are released into or consumed from the environment. We ask what determines invasion outcomes in such microbial communities. Here, we use a model based on chemical-mediated interactions among microbial species to assess the impact of positive and negative interactions on invasion outcomes. We classified invasion outcomes as resistance, augmentation, displacement, or disruption depending on whether the richness of the resident community was maintained or dropped and whether the invader was maintained in the community or went extinct. We found that as the number of invaders increased relative to size of the resident community, resident communities were increasingly disrupted. As facilitation of the invader by the resident community increased, resistance outcomes were replaced by displacement and augmentation. By contrast, as facilitation increased among residents, displacement outcomes shifted to resistance. When facilitation of the resident community by the invader was eliminated, augmentation outcomes were replaced by displacement outcomes, while when inhibition of residents by invaders was eliminated, there was little change in the frequency of invasion outcomes. These results suggest that a better understanding of the chemical-mediated interactions within resident communities and between residents and invaders is crucial to predicting the success of invasions into microbial communities.

## Introduction

Members of resident communities can impact the success or failure of invading species to establish in an ecosystem. Residents can affect invasion outcomes by altering resource availability, occupying available niches, or interacting with them directly or indirectly via predation, competition, facilitation, or other mechanisms. The relative importance of these factors in determining invasion outcomes varies between communities and ecosystems. Functional composition of resident communities, for example, is an important determinant of invasion success in many grasslands (1), while release from consumer or competitive pressure is an especially important factor in marine invasions (2). Among microbes, interactions often occur via chemical mediators released into the environment (3,4). These mediated interactions are believed to be influential in many microbial communities, but their importance to invasion outcomes remains unexplored.

In human microbiota, where preventing invasion is a first step in preventing many diseases, this phenomenon is sometimes referred to as colonization resistance. The potential for resident microbes to protect us from pathogens has been observed as early as 1917, by the discovery of *Escherichia coli* Nissle 1917 that antagonized and blocked enteric pathogens (5). More examples across different microbiota sites abound: nasal microbiota protects us against respiratory *Staphylococcus aureus* infection (6,7), gut microbiota protects us from *Clostridium difficile* infection (8,9), and oral microbiota can block *E. coli* infection (10), to name a few. While previous research shows that resident microbiota can suppress pathogens (11–13), the mechanism of colonization resistance and the role of interspecific interactions is not fully understood (10).

The majority of explanations about how microbiota achieve colonization resistance is built around competition (14). This competition can take many forms (15), including resource competition, production of toxic compounds, or induction of host immunity. Positive interactions and their role, especially within the context of a community interacting via diffusing chemical mediators, have been hardly investigated. This is despite the fact that in invasion ecology (16–19), a topic closely linked to colonization resistance, the role of facilitation—interactions that benefit other community members—has been considered for decades (20,21). To highlight a few examples, it has been shown that incidental mutual interactions with native species promotes invasion (22), facilitation among two non-native species can make invasion more successful (23), and facilitative ecosystem-engineering can increase invasion success (20).

Strategies to prevent or assist the addition of new taxa to human microbiota can be critical components of healthcare. For example, current approaches to promoting the development of a healthy neonatal gut microbiome include interventions in the maternal microbiota during pregnancy, dietary changes, exploration of environmental impacts, and application of prebiotics and probiotics. Exploration of the community and environmental contexts in which each strategy is likely to be effective could benefit from guidance from appropriate theory. However, with a few exceptions (such as (10)), the existing theoretical mechanisms of colonization resistance and invasion ecology have not fully made their way into microbiota intervention strategies. This despite the fact that several past efforts on invasion and colonization resistance were performed in microbial systems (24–27).

Mechanistic studies typically fall along a spectrum of generality. In invasion ecology, on one side are general theoretical predictions (19,21,28), while on the other side are specific, although remarkable, instances such as host-supported colonization of legume by rhizobia (29). Studies in human microbiota can be done in these two types as well (see a summary of different model systems in (30), for example). Here, we choose to focus on developing general insights that could inform and guide future microbiota-based intervention strategies. Since such general insights are hard to draw in natural microbiota, where members and interactions are not often adequately known, we use *in silico* models in our approach instead. We use a previously introduced mathematical model of microbes and explicit mediators of interactions (31,32). Using this model, we investigate how invasion of microbial communities is affected by chemical-mediated interspecific interactions between the invader and resident members or among resident members. Our results suggest that interspecies interactions can markedly influence colonization resistance.

## Results

### Increasing the propagule size does not increase the chance of incorporation of an invader into a resident community

To assess colonization resistance, we set up an *in silico* invasion assay in which stable *resident* communities are challenged with invaders introduced at different fractions (Fig 1A; see Methods for details). We categorize the outcome based on the fate of the invader and the resident community. There are four possible outcomes (Fig 1A): ‘Resistance’ (invader extinct, all resident species maintained), ‘Augmentation’ (invader maintained, all resident species maintained), ‘Disruption’ (invader extinct and some resident species also go extinct), and ‘Displacement’ (invader maintained and some resident species go extinct). We observe that only when the relative size of the invader population introduced into the resident community—hereafter called propagule size—is comparable to the resident community, the chance of observing different outcomes is affected (Fig 1B and S1). In our model, no considerable change in outcomes is observed below a propagule size of ~10%. Community outcomes are affected when invader propagule size exceeds ~10% of the resident population. Probability of resistance decreases across this range. Importantly, in our model, a larger propagule size does not appear to alter the chance of augmentation. At large propagule sizes, the invader is maintained in the final population only at the cost of losing some of the resident species. This is at odds with common wisdom of using probiotic at high doses to allow the “helpful” microbes to be augmented into the microbiota.

**Fig 1.**
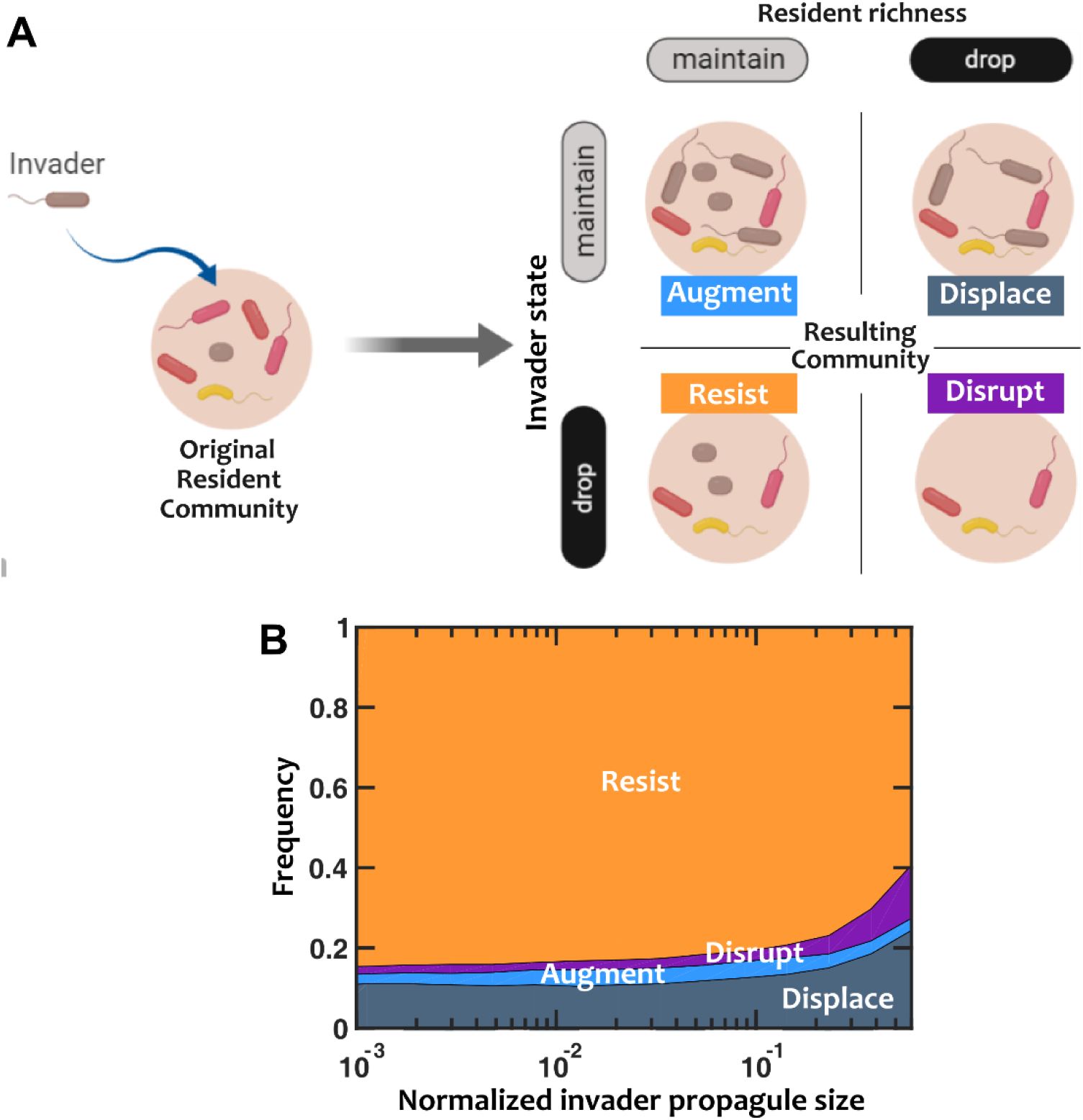
Only at large invader propagules colonization resistance is weakened. (A) In our *in silico* invasion assay, we first assemble instances of stable resident communities and then assess the outcomes after introducing an invader. Based on whether the invader persists or goes extinct and whether the resident community maintains or decreases in richness, we categorize the outcomes into four groups: ‘Resistance’ (invader extinct, resident species maintained), ‘Augmentation’ (invader maintained, all resident richness maintained), ‘Disruption’ (invader extinct, resident richness drops), and ‘Displacement’ (invader maintained, resident richness drops). (B) As the normalized propagule size (*i.e.* the amount of invader cells introduced, relative to the total population size of the resident community) increases, the probability of resistance decreases, the probability of disruption and displacement increases, and the probability of augmentation remains approximately constant. Number of instances examined N_s_=1000. Interactions among resident members are equally likely to be facilitative or inhibitory (*f*_*fac*_=0.5). Interactions between resident members and the invader are mostly inhibitory (*f*_*fac,inv*_=0.1). The invader has on average a 50% advantage in basal growth rate compared to resident members (*r*_*0,inv*_/*r*_*0,res*_=1.5). To visualize the trends more clearly, here we do not include the error-bars (see Fig S1 for confidence intervals).

### Invaders with higher basal growth rate are more likely to displace residents

We asked if the basal growth of the invader—what would conventionally be the main indicator of its competitive potential—is the major determinant of invasion success. To answer this question, we introduced invaders with different basal growth rate—i.e. growth rate in the absence of interactions—into the community and tallied the invasion outcomes (Fig S2). The results show that as the community is challenged with invaders with basal growth rates higher than resident members, the outcome shifts from resistance to displacement. The augmentation and disruption outcomes remain unlikely as the basal growth rate of the invader increases.

### Facilitation of invader by resident microbiota can weaken colonization resistance

To investigate the impact of interactions on invasion outcomes, we first looked at the interactions between the resident microbiota and the invader. We kept the interactions within the resident communities fixed and transitioned the interactions imposed by the resident microbiota on the invader from mostly inhibition to mostly facilitation (Fig 2). The results show a clear trend: when the resident community facilitates the invader, the chance of invasion (augmentation or displacement outcomes) is enhanced. As facilitation of the invader increases, the fraction of resistance decreases, with a corresponding increase in displacement. The chance of augmentation increases slightly, while the probability of disruption remains relatively constant.

**Fig 2.**
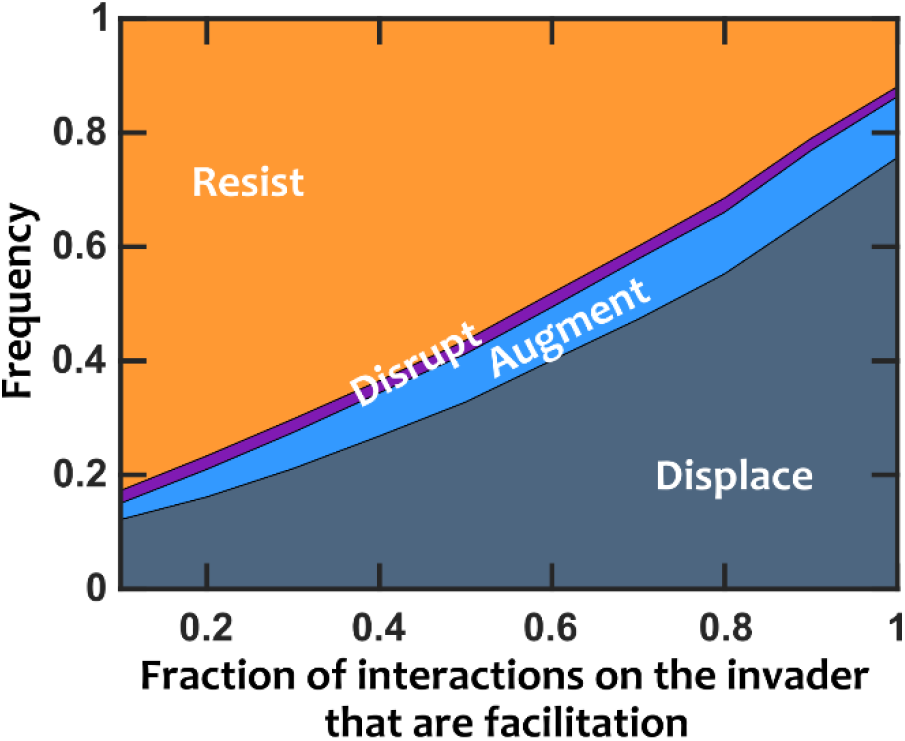
When resident species facilitate the invader, colonization resistance is weakened. Invasion success drastically increases when we switch the interactions that influence the invader from inhibition to facilitation. Number of instances examined N_s_=1000. Interactions among resident members are equally likely to be facilitative or inhibitory (*f*_*fac*_=0.5). Normalized basal growth rate of the invader is 1.5. Normalized introduced propagule size is 0.3%.

This result is fairly intuitive. If the resident members mostly facilitate the invader, the growth rate boost that the invader receives makes the invasion more successful. The prevalence of the displacement category in these results rather than augmentation (i.e. successful invasions tend to deplete resident community richness) reinforces the view that facilitation of invaders may be detrimental to resident communities.

### More cooperative microbiota show stronger colonization resistance

We shifted our focus to the interactions within the resident microbiota to assess their impact on invasion outcomes. For this, we surveyed many examples of stable resident communities formed by groups of species that engaged in interactions at different facilitation to inhibition fractions. The results showed that invasion was less successful when there was prevalent facilitation among resident members (Fig 3). This trend holds when the resident members are mostly antagonistic against the invader as well (Fig S3).

**Fig 3.**
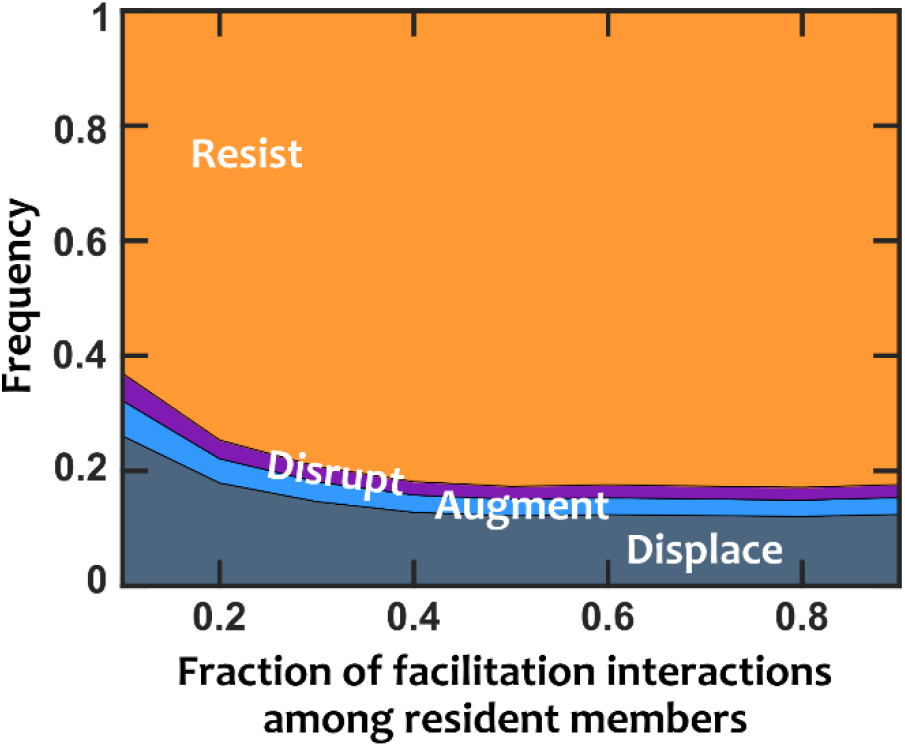
Facilitation among resident species strengthens colonization resistance. Invasion success decreases when interactions among resident species are predominantly facilitation rather than inhibition. The interactions between resident species and the invader are mostly inhibitory (*f*_*fac,inv*_=0.1). Normalized basal growth rate of the invader is 1.5 (compared to resident members). Normalized introduced propagule size is 0.3%.

One explanation for the above results is from the perspective of available interactions. The mediators present in the resident community can modulate the growth rate of the invader positively or negatively. If other members of the community are already mostly positively benefitting from these mediators, that leaves less room for an invader to take advantage of those resources to keep up with resident members or outpace them in growth.

### The net interaction between the invader and resident microbiota determines the type and strength of colonization resistance

What are the mechanisms through which interactions between the invader and resident members determine the chance of invasion? We considered a particular case where the chemical mediators of the resident were more likely to be inhibitory to the invader. We asked how much this inhibitory effect as well as consumption or production of mediators by the invader influenced the invasion outcome. The invader interfaces with residents in two ways: it can affect the chemical environment of the resident community by consuming or producing chemical mediators, and it also gets affected by the chemicals in the environment. We tested how removing production, consumption, and/or chemical influence on the invader impacted invasion outcomes (Fig 4 and Fig S5).

**Fig 4.**
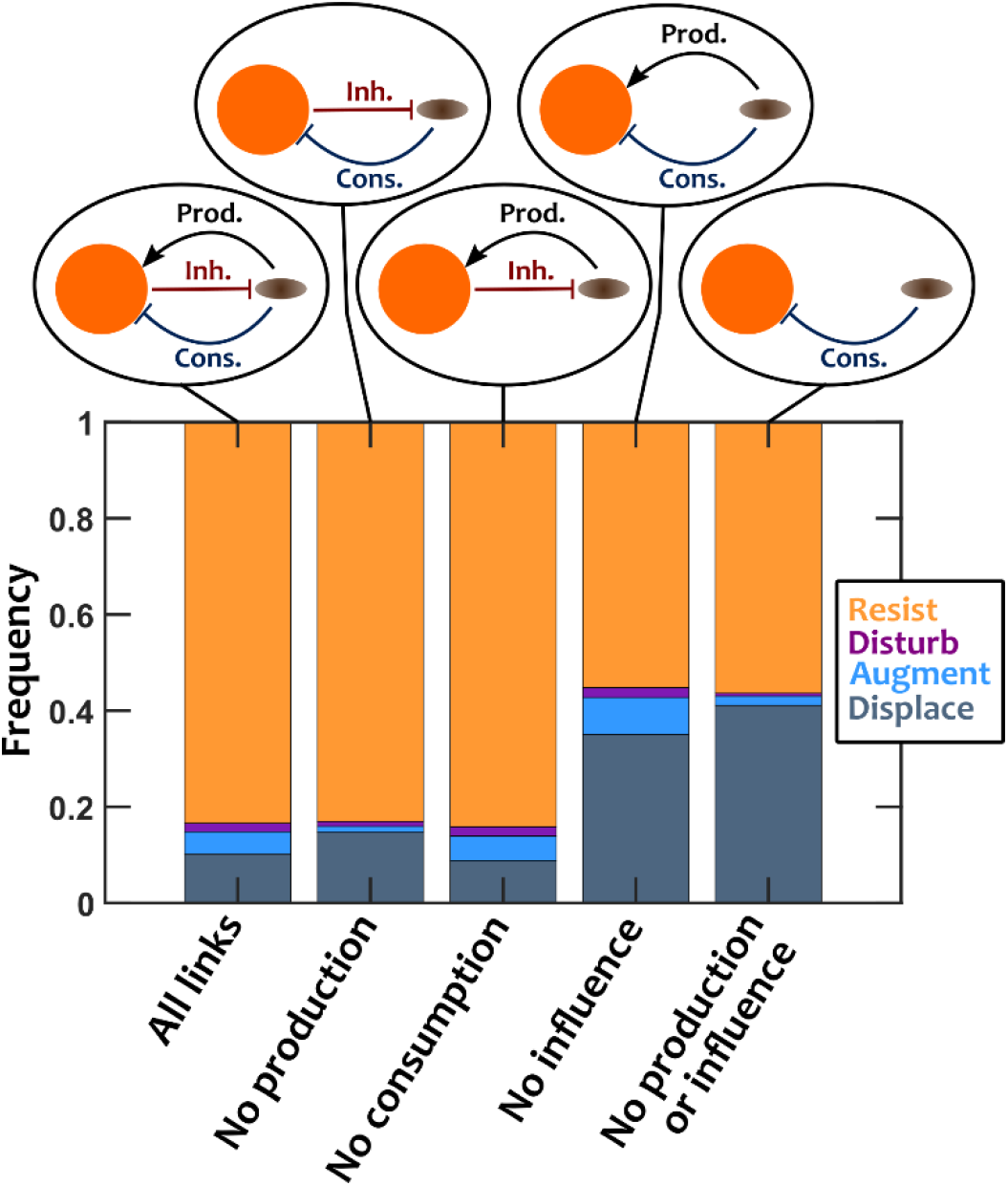
The effective interaction between the resident community and the invader alters the outcome of invasion. We examined how invasion outcomes changed when the interactions between the invader and resident members were altered. In the original case, the invader is inhibited by resident species, and the invader consumes and produces some chemical mediators. In other cases, the invader inhibition, consumption, or production (or a combination of them) are removed from the system, as shown in the insets, to assess the impact of each. Interactions among resident species are equally likely to be facilitative or inhibitory (*f*_*fac*_=0.5). The influence of residents on the invader is mostly inhibitory (*f*_*fac,inv*_=0.1). Invader has a normalized basal growth rate of 1.5. Number of instances examined N_s_=10000.

To interpret the results, we first recall that the resident community involves mostly facilitative interactions among resident members (32). Thus, we expect production of mediators by the invader to benefit the resident community, while consumption of mediators will inhibit residents. We can simplify our view of the system as two compartments—*i.e.* the resident community and the invader—and their interactions. As shown in the left-most inset of Fig 4, in the original case the invader facilitates the resident community through production of mediators, inhibits the resident community through consumption of mediators, and gets inhibited by the chemical environment of the resident community (our assumption for this example). Examining the results after removing different components of this model leads to several interesting observations: (1) Removing invader production (thus, no facilitation) removes the augmentation types from the outcomes and increases the chance of displacement. (2) Removing consumption (thus, no inhibition of microbiota by the invader) only marginally decreases the chance of displacement and slightly improves colonization resistance. (3) Removing the inhibition of the invader increases the chance of invasion, primarily by contributing to the displacement outcomes. The results are overall in agreement with what we would expect from a two-component system (33): facilitation increases stable coexistence (manifested as the augmentation outcome), whereas mutual inhibition increases bistability (most outcomes are resistance or displacement).

Insights from the impact of different interactions on invasion and colonization resistance outcomes (Fig 4 and Fig S5) is a helpful first-step to rethink the intervention strategies for manipulating microbiota. Our results suggest that when interactions other than pure competition are involved, propagule size is not an effective parameter to influence the outcomes (Fig 1). Instead, interactions between the invader and the resident community and the interactions within the resident community appear to influence the outcome (Fig 4). Take a situation of assimilating a probiotic strain into an existing microbiota. Our results suggest that rather than administering larger doses of the probiotic strain, the focus should be on creating facilitation or removing existing inhibition between the probiotic strain and the resident microbiota.

## Discussion

To examine how interspecific interactions affect invasion outcomes within microbiota, we used a mediator-explicit model of chemical interactions among microbes (31,32). We developed an *in silico* yet empirically feasible assay to assess different types of invasion. Invasion outcomes in this assay are categorized as resistance, augmentation, disruption, or displacement depending on whether the invader is maintained in the community or driven extinct and whether the resident community maintains its richness or some resident species go extinct. Using this model, we investigated the impact of different parameters—including those related to interactions—on invasion outcomes. We saw that as the size of the invader population increased relative to the resident community, resistant outcomes decreased and were replaced by disruptions. With invaders that higher basal growth rates relative to the resident members, resident communities were increasingly disrupted. As expected, we found that invaders are more successful if the chemical environment created by the resident community facilitates them. We also observed that if the interactions among members that form the resident community are more facilitative—rather than inhibitory—colonization resistance is strengthened. Finally, we found that when facilitation of the residents by the invader is removed, augmentation outcomes are replaced by displacement, while when inhibition of residents by invaders is removed, there is little change in invasion outcomes.

The tradeoff in prevalence between resistance and displacement outcomes—created by facilitative and inhibitory interspecific interactions, respectively—in our results already highlights how such interactions influence colonization resistance. This on its own is not a new finding. Many studies before us have pointed to the impact of interactions, positive or negative, on invasion, including (10,14,17,20) to name a few. Our contribution is to highlight this impact in a model that resembles continuous growth in microbiota. We propose a framework based on consumption and production of chemicals that can impact cell growth and track down how these processes lead to an eventual success or failure of an invader. It is striking to note the many different ways invasion trajectories can be altered by modifying or removing interspecific interactions. An increase of facilitation of the invader, for example, leads to an increase in displacement and augmentation, both outcomes in which the invader is maintained in the final community. By contrast, facilitation among residents increases colonization resistance, while the loss of facilitation of residents by the invader leads to a shift from augmentation, in which the full resident community richness is maintained, to displacement, in which one or more residents are driven extinct. The context dependence of the effects of positive interactions on invasion outcomes emphasizes the importance of improving our understanding of the natural history of microbial communities.

A simple two-compartment model of invader versus the resident community offers an intuitive prediction of invasion outcomes. If the effective interaction between the resident community and the invader is mutual inhibition, we would expect bistability—either the resident community resists invasion or it crumbles. In contrast, for assimilation of an invader into microbiota, a facilitative interaction between the resident members and the invader appears necessary. When interactions are the main driving force, two-way facilitation in a mutualism or one-way facilitation in a commensalism or prey-predation interaction are necessary for stable coexistence of the two components (33).

This work was conducted using a model in which microbes interact exclusively via chemical mediators released into the environment. Such interactions are common in microbial communities and are important determinants of community assembly and other processes, but they are by no means the only important microbial interactions. For example, in many microbiota, contact dependent growth inhibition (34) and type VI secretion systems (35) are critical mechanisms of bacterial competition that rely on contact between interacting cells. Many interactions are also mediated by induction of the host immune system (36). This work cannot account for the possible roles of such interactions influencing invasion trajectories or predict the relative importance of chemical-mediated interactions on invasion outcomes. Direct comparison of the impact of these alternative interaction mechanisms on invasion outcomes would provide an important extension of this work.

In our model, we intentionally have focused on interspecies interactions beyond competition for resources. An important question is whether the trends we have found will hold when resource competition is included in the mode. To answer this question, we modified our model and explicitly incorporated competition for a single limiting resource (Materials and Methods). We chose the amount of resource such that within each round of growth the populations would deplete the resource, ensuring that resource competition was in effect. We found that making the resource limitation more stringent did not considerably influence the outcome of invasion (Fig S7). Our results show that the low sensitivity of outcomes to the propagule size still holds in this modified model (Fig S8). Changing the basal growth rate of the invader also resulted in an overall trend resembling the case with no explicit resource competition, with only one qualitative difference: an increased occurrence of disruption at intermediate levels of invader growth rate (Fig S9). The trends of invasion outcomes based on invader-resident interactions (Fig S10) and resident-resident interactions (Fig S11) also largely remained intact.

Finally, by helping us to understand under what circumstances particular invasion outcomes are likely, this work can help to guide interventions towards those that are appropriate for restructuring microbiota. For example, the common strategy of increasing the dosage of probiotics to increase the chance of invasion is unlikely to succeed, because increasing the invader propagule size has little effect on invasion outcomes, except when the invader is almost as prevalent as the entirety of the resident community and even then disruption increases rather than augmentation, which is typically the desired outcome of such interventions. Each of the four invasion outcomes described here has a real-world equivalent in the human microbiota in which it is the desired state or outcome of an intervention. For example, resistance is typically the desired outcome of invasion into a healthy microbiota. There are many microbial communities that are known to resist invasion by pathogens, such as (5–9). Our work suggests that interactions are likely a key component of this resistance, which would be favored in communities that inhibit the particular invader in question and have many facilitative interactions among residents. This implies that predicting, discovering, and improving upon successful microbiota interventions would be aided by a deeper, more species-specific understanding of the interactions that operate within microbial communities.

## Materials and Methods

### Mathematical model

We use a mediator-explicit model that includes the species and the chemical environment that mediates interspecies interactions.

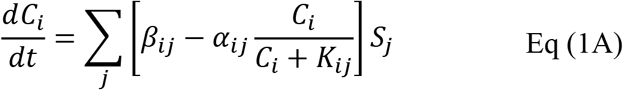

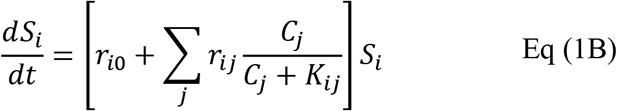

In which *C*_*i*_ and *S*_*i*_ are the concentrations of the mediators and cell densities, respectively; *β*_*ij*_ and *α*_*ij*_ are production and consumption rates, respectively; and *K*_*ij*_ is the saturation concentration for uptake and growth rate influence. The following parameters are used throughout this work: *r*_*i*0_ has a uniform distribution between 0.08 and 0.12 hr^−1^; *r*_*ij*_’s amplitude has a uniform distribution between 0 and 0.2 hr^−1^, and its sign can be positive or negative with a probability that is specified in each case; *β*_*ij*_ has a uniform distribution between 0.05 and 0.15 fmole/cell per hour; *α*_*ij*_ has a uniform distribution between 0.25 and 0.75 fmole/cell per hour; and *K*_*ij*_ has a uniform distribution between 50 and 150 nM. The initial cell density is 10^4^ cells/ml, and the community is cyclically diluted back to this initial value when the density reaches 10^7^ cells/ml. This dilution scheme—replicating conventional growth situation in the lab—ensures that most shared resources are replenished during community growth (32).

The model with explicit resource competition is different from the basic model in that it also includes a single resource that all species require for growth.

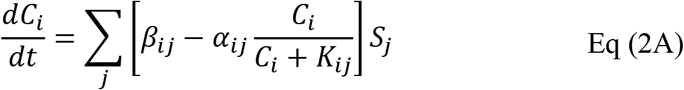

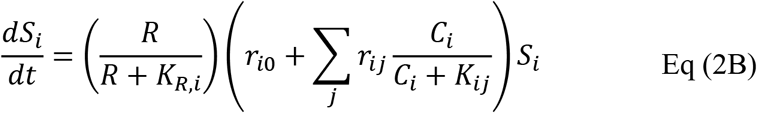

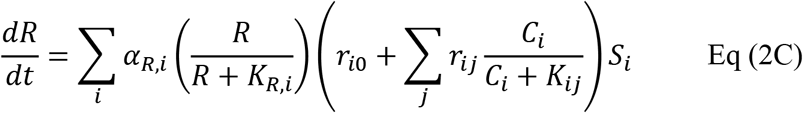

We have assumed that the consumption of resource *R* is directly proportional to the growth rate of species, with a given resource consumption rate *α*_*R,i*_. Resource *R* also affects the growth rate in a saturating fashion, with half-maximum concentration of *K*_*R,i*_. Unlike other mediators, the resource *R* is not produced by any of the species; it is supplied in the medium and thus replenished at dilution steps. For these simulations, dilution either happens when the density reaches 10^7^ cells/ml or after 80 hours. This choice is made to allow population turn-over even when the amount of resources in the environment is low to the level that the density does not reach the dilution threshold. The number of dilution cycles are adjusted in each case such that the culture experiences a total of 200 generations (regardless of the amount of resources available) to allow a fair comparison.

Equations (with or without an explicit resource) are numerically solved using a forward Euler method. We chose the time-step for simulations to be at least ten times smaller than the shortest doubling time in each case. This choice offers a trade-off between accuracy and total simulation time. We have tested smaller time-steps and confirmed that the outcomes were not affected.

### Parameters used for simulations

Unless specified, the following parameters were used in simulations in this manuscript

| Parameter | Description | Value |
| --- | --- | --- |
| $N_c$ | Number of different species in the initial pool used for obtaining instances of resident communities | 20 |
| $N_m$ | Number of chemical mediators in the simulations that mediate the interactions among species | 10 |
| $N_s$ | Number of instances of simulations run for each case | 1000 |
| $N_{gen}$ | Number of generations simulated to obtain stable resident communities; also the number of generations simulated to assess invasion success | 200 |
| $F_{inv}$ | Range of invader introduction fractions examined for assessing invasion (20 points, equally spaced in log scale) | $10^{-4}$ to 0.99 |
| $N_{inoc}$ to $N_{dil}$ | Range of total population density simulated (from inoculation to dilution threshold) | $10^4$ to $10^7$ cells/ml |
| $N_{ext}$ | Extinction population density per species | 0.1 cells/ml |
| $K_{sat}$ | Average chemical concentration threshold for saturation of chemical consumption and fitness influences (average of $K_{ij}$ values) | $10^4$ μM |
| $r_0$ | Basal net growth rate of community members, in the absence of interactions (uniform random distribution) | 0.08-0.12 $\text{hr}^{-1}$ |
| $\alpha_{ij}$ | Consumption rate of chemicals by species, per cell (uniform random distribution) | 0.25-0.75 fmole.$\text{hr}^{-1}$ |
| $\beta_{ij}$ | Production rate of chemicals by species, per cell (uniform random distribution) | 0.05-0.15 fmole.$\text{hr}^{-1}$ |
| $r_{i0}$ | Amplitude of the influence of chemicals on species' growth rate (uniform random distribution) | 0-0.2 $\text{hr}^{-1}$ |
| $f_{fac}$ | Fraction of non-neutral interactions among the initial pool of resident microbes that are facilitative | 0.5 |
| $r_{i0,inv}$ | Amplitude of the influence of chemicals on invader's growth rate (uniform random distribution) | 0-0.2 hr<sup>-1</sup> |
| $f_{fac,inv}$ | Fraction of non-neutral interactions affecting the invader that are facilitative | 0.5 |
| $q_p$ | The degree of connectivity of producer species to chemicals both for the resident members and the invader (forming a binomial network) | 0.3 |
| $q_c$ | The degree of connectivity of chemicals to species that they influence both for the resident members and the invader (forming a binomial network) | 0.3 |
| $R_0$ | Concentration of the limiting resource in fresh medium | 10<sup>6</sup> fmole/ml |
| $K_R$ | Average saturation concentration of the limiting resource | 1 mM |
| $\alpha_R$ | Consumption rate of limiting resource, per cell (uniform random distribution) | 5-15 fmole.hr<sup>-1</sup> |

### *In silico* invasion assay

To assess invasion success, we first assemble instances of stable resident communities *in silico*, following the enrichment procedure outlined in (32). For each instance, we put together 20 species that are interacting through 10 chemical mediators. We simulate the dynamics in growth-dilution cycles for 200 generations to identify instances of stable *in silico* communities. We then introduce invaders at an initial fraction of 0.3%. We show that this fraction is a representative of the outcome at “small” propagule sizes (see Fig 1). We simulate the dynamics in growth-dilution cycles for an additional 200 generations. At the end of these simulations, we calculate the fraction of the invader cells in the resulting community. Based on the relative fraction of the invader between the initial and resulting communities across different propagule sizes, we categorize the invasion outcome as resistance, augmentation, displacement, or disruption. Resistance: resident community richness is maintained, invader population drops. Augmentation: resident community richness is maintained, invader population is maintained. Displacement: resident community richness drops, invader population is maintained. Disruption: resident community richness drops, invader population drops. To obtain reliable statistics about invasion outcomes, we repeat the process of assembling resident communities and challenging them with invaders for at least 1000 times.

## Acknowledgements

This work was supported through a start-up fund by Boston College and through an Award for Excellence in Biomedical Research by the Richard and Susan Smith Family Foundation. Figure 1A was created with BioRender.com.

## Supplementary Information

**Fig S1.**
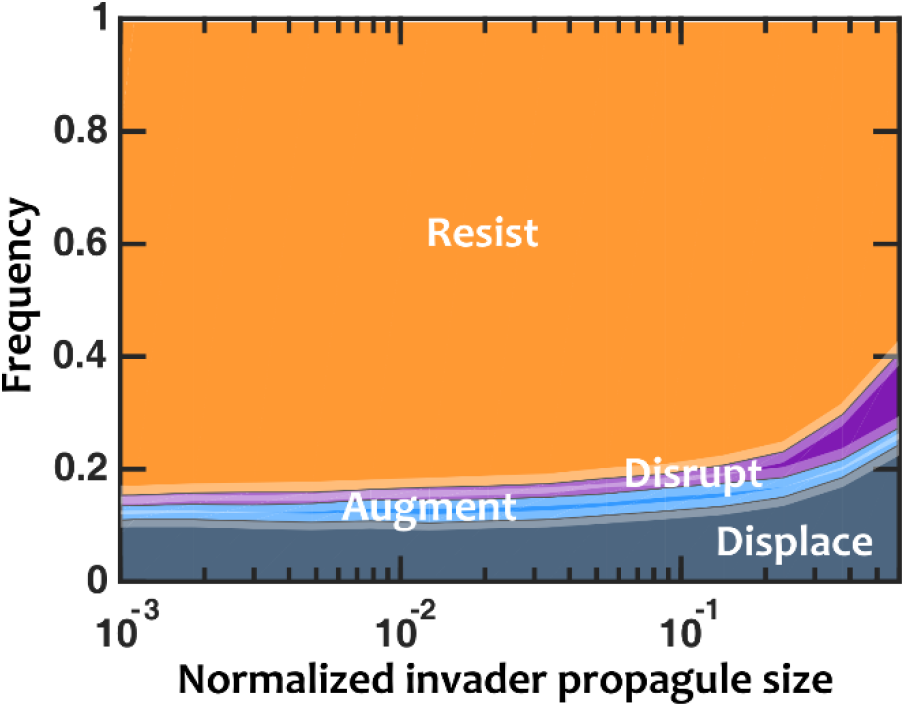
Only at large invader propagules colonization resistance is weakened. Here we have added confidence intervals to the graph, with all the parameters being similar to Fig 1B. Confidence intervals for outcome frequencies are calculated using the Clopper-Pearson method (using binofit function in Matlab). 80% confidence intervals are plotted as a shaded region around each mean frequency. Number of instances examined: N_s_=1000.

**Fig S2.**
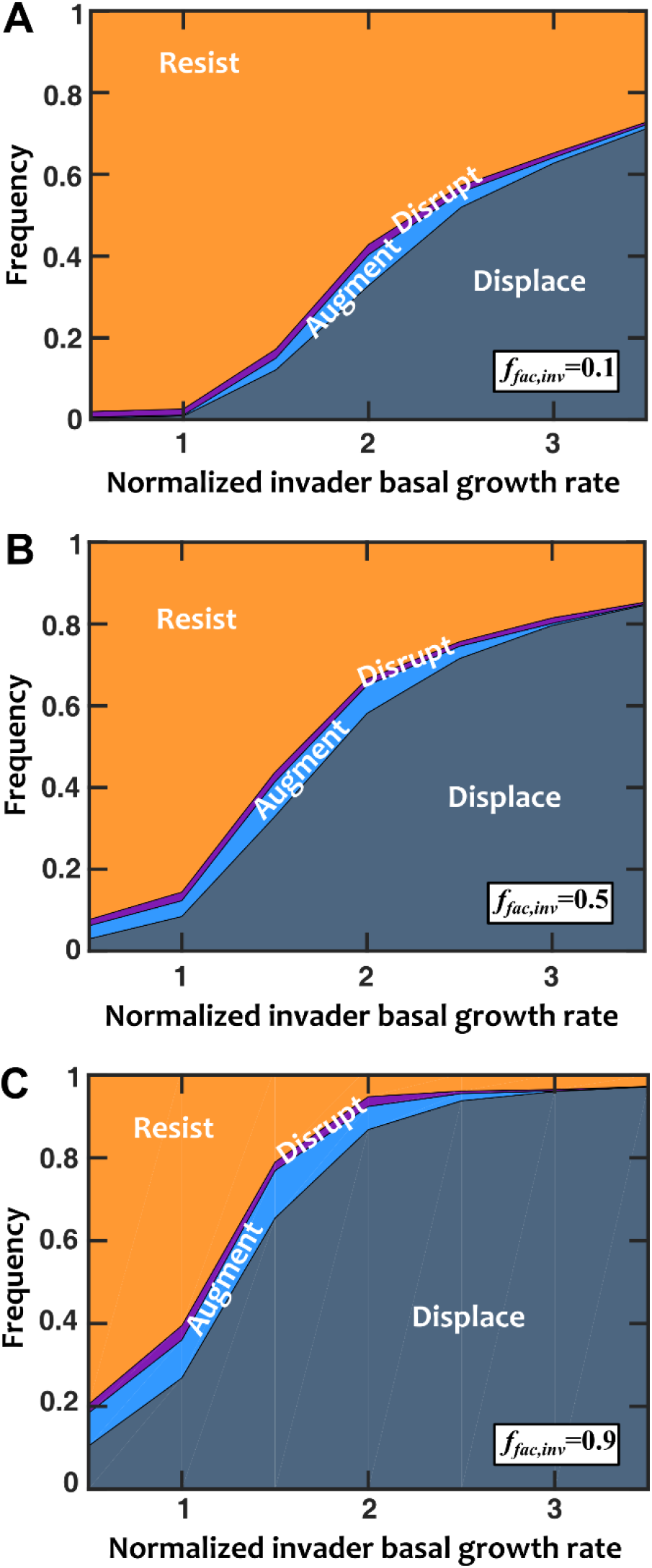
Invaders with a higher basal growth rate shift invasion outcomes primarily from resistance to displacement. The pattern holds when the influence of mediators on the invader is (A) mostly inhibitory (*f*_*fac,inv*_=0.1), (B) equally facilitative or inhibitory (*f*_*fac,inv*_=0.5), or (C) mostly facilitative (*f*_*fac,inv*_=0.9). In all cases interactions among resident species are equally likely to be facilitative or inhibitory (*f*_*fac*_=0.5). Normalized basal growth rate of the invader is relative to resident species. Number of instances examined N_s_=1000.

**Fig S3.**
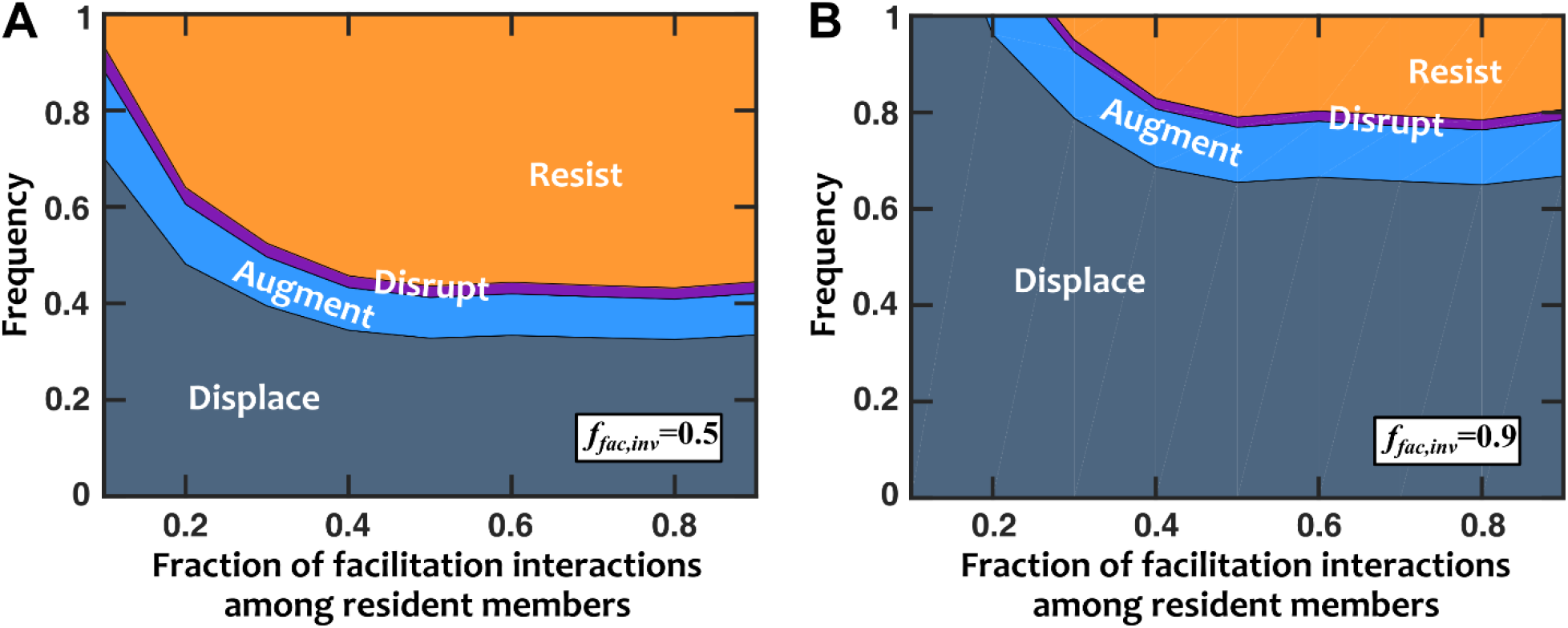
Facilitation among resident species strengthens colonization resistance. Invasion success decreases when interactions among resident members are predominantly facilitation rather than inhibition. The interactions of resident species with the invader are (A) equally likely to be facilitative or inhibitory (*f*_*fac,inv*_=0.5) or (B) mostly facilitative (*f*_*fac,inv*_=0.9). In all cases interactions among resident species are equally likely to be facilitative or inhibitory (*f*_*fac*_=0.5). Normalized basal growth rate of the invader is 1.5 (compared to resident members). Number of instances examined N_s_=1000.

**Fig S4.**
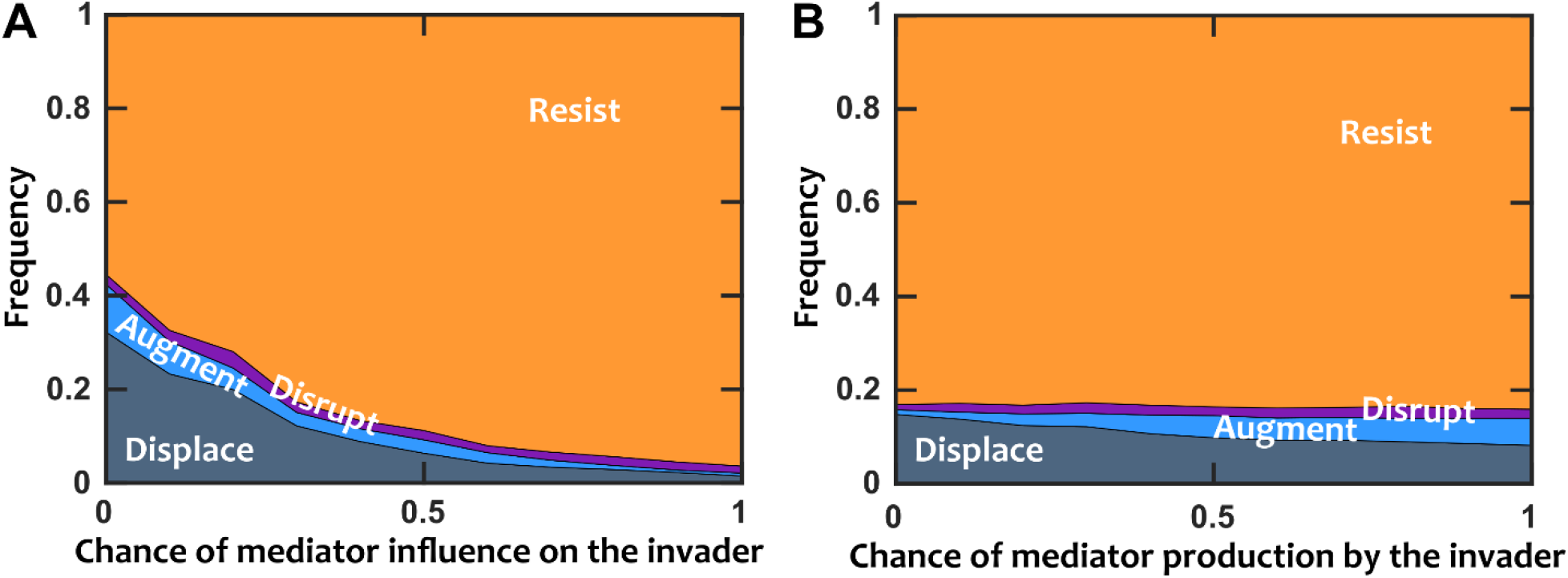
Invader connectivity affects invasion outcomes. (A) When the chance of mediators influencing the invader increases, colonization resistance is strengthened, as expected. (B) When the chance of the invader producing chemical mediators increases, augmentation becomes more likely. Interactions between resident species and the invader are mostly inhibitory (*f*_*fac,inv*_=0.1). Interactions among resident species are equally likely to be facilitative or inhibitory (*f*_*fac*_=0.5). Normalized basal growth rate of the invader is 1.5 (compared to resident members). Number of instances examined N_s_=1000.

**Fig S5.**
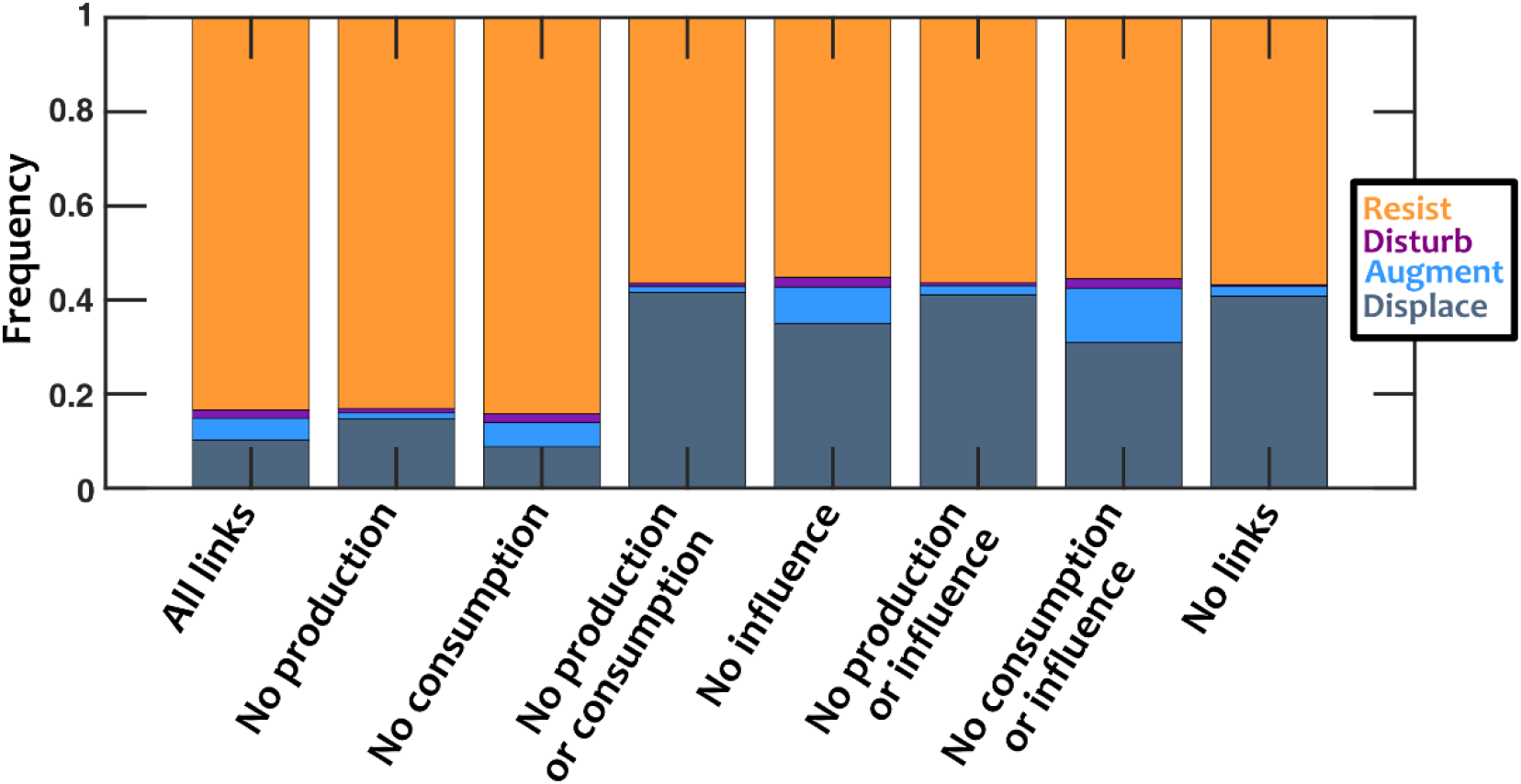
The effective interaction between the resident community and the invader alters the outcome of invasion. We expanded the results in Fig 4 (all parameters kept the same) to demonstrate all eight possible combinations of keeping or removing production, consumption, or mediator influence. Interactions among resident species are equally likely to be facilitative or inhibitory (*f*_*fac*_=0.5). The influence of residents on the invader is mostly inhibitory (*f*_*fac,inv*_=0.1). Invader has a normalized basal growth rate of 1.5. Number of instances examined N_s_=10000.

**Fig S6.**
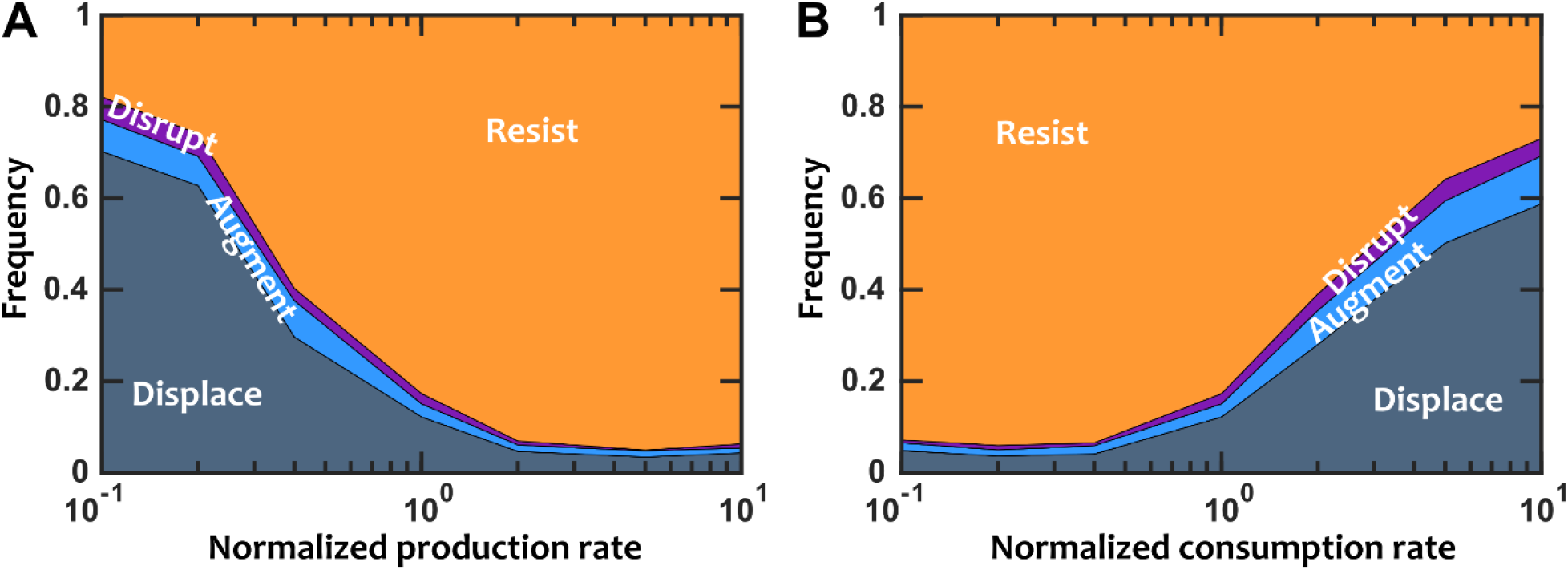
Colonization resistance is weakened when the mediators that inhibit the invader are depleted. (A) When the production of mediators increases, colonization resistance is strengthened. (B) When the consumption of the mediators is increased, mediators are depleted and thus colonization resistance is weakened. Interactions between resident species and the invader are mostly inhibitory (*f*_*fac,inv*_=0.1). Interactions among resident species are equally likely to be facilitative or inhibitory (*f*_*fac*_=0.5). Normalized basal growth rate of the invader is 1.5 (compared to resident members). Number of instances examined N_s_=1000.

**Fig S7.**
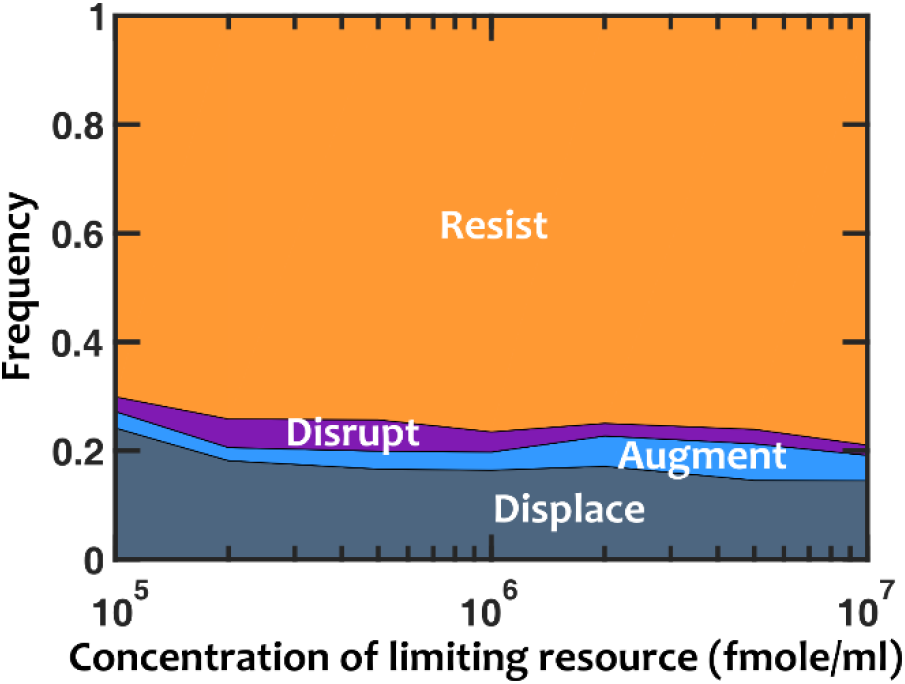
With explicit resource competition, the extent of resource limitation only marginally influences the outcomes. Interactions between resident species and the invader are mostly inhibitory (*f*_*fac,inv*_=0.1). Interactions among resident species are equally likely to be facilitative or inhibitory (*f*_*fac*_=0.5). Normalized basal growth rate of the invader is 1.5 (compared to resident members). The invader is introduced at 0.3% of the resident community size. The amount of the limiting resource is varied between 10^5^ and 10^7^ fmole/ml. Each invasion assay is run for as many dilution rounds as needed to reach 200 generations of total community growth. Number of instances examined N_s_=1000.

**Fig S8.**
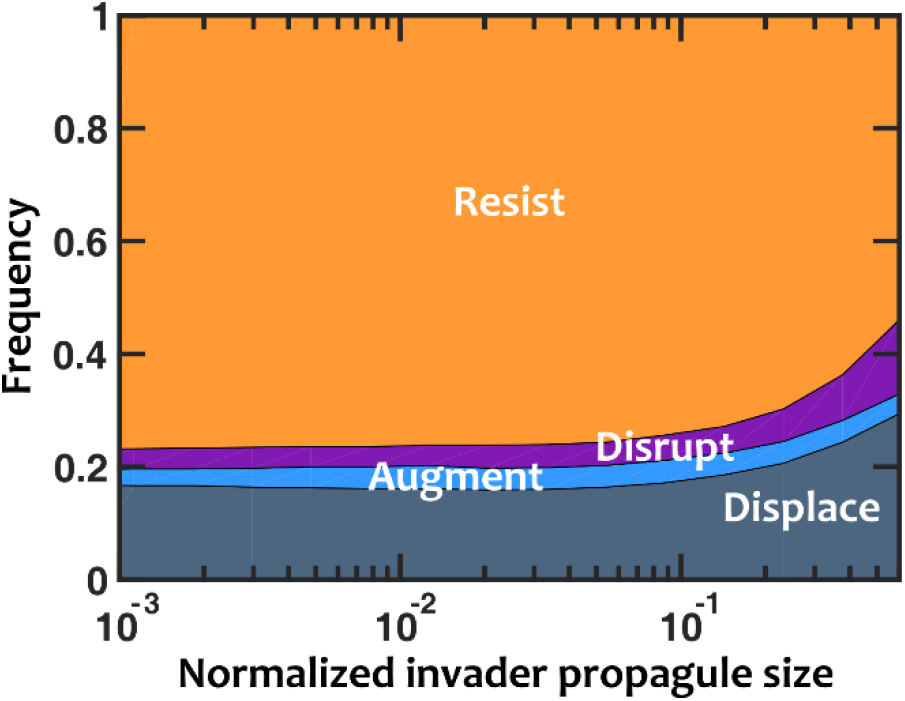
Only invader propagules comparable to community size can weaken colonization resistance, when explicit resource competition in included. As the normalized propagule size increases, the probability of resistance decreases, the probability of disruption increases, and the probability of augmentation or displacement remains approximately constant. Number of instances examined N_s_=1000. Interactions among resident members are equally likely to be facilitative or inhibitory (*f*_*fac*_=0.5). Interactions between resident members and the invader are mostly inhibitory (*f*_*fac,inv*_=0.1). *r*_*0,inv*_/*r*_*0,res*_=1.5. The amount of the limiting resource is set at 10^6^ fmole/ml.

**Fig S9.**
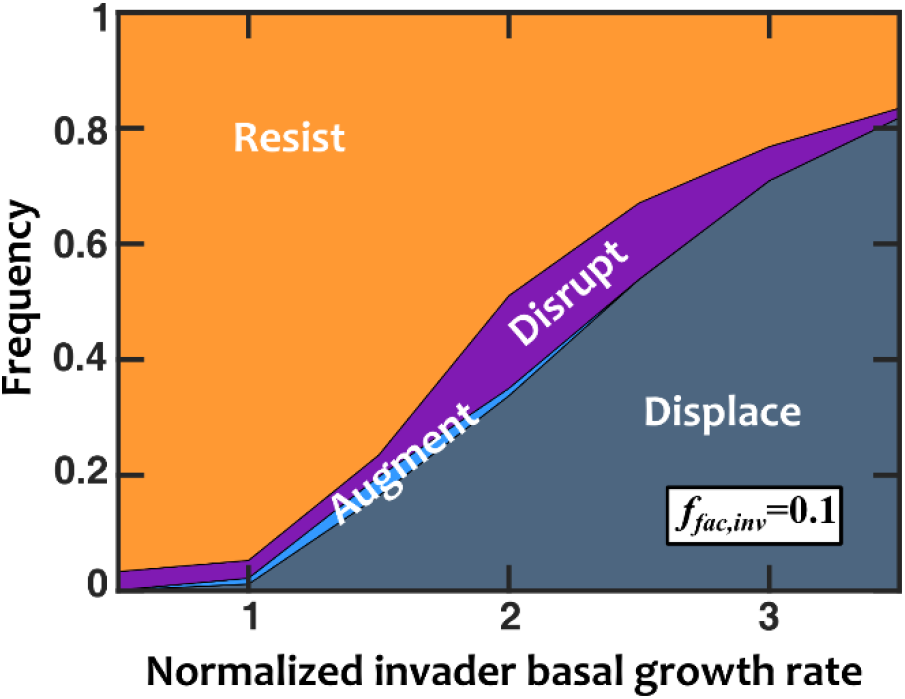
Invaders with a higher basal growth rate shift invasion outcomes primarily from resistance to displacement, when a single limiting resource is added to the model. The influence of mediators on the invader is mostly inhibitory (*f*_*fac,inv*_=0.1). Interactions among resident species are equally likely to be facilitative or inhibitory (*f*_*fac*_=0.5). Normalized basal growth rate of the invader is relative to resident species. Number of instances examined N_s_=1000. The amount of the limiting resource is set at 10^6^ fmole/ml.

**Fig S10.**
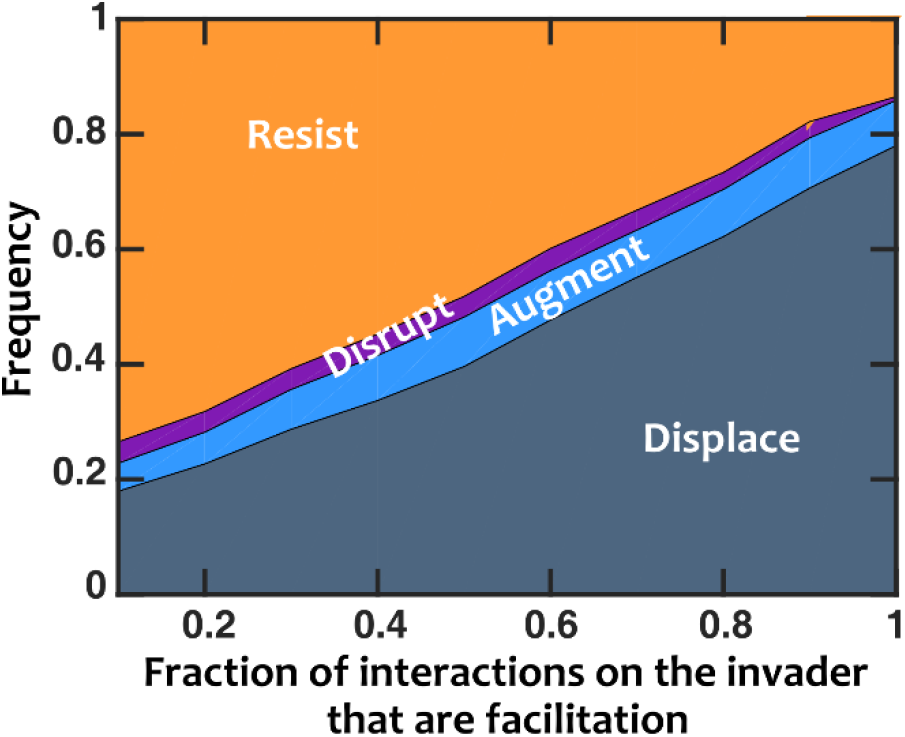
When resident species facilitate the invader, colonization resistance is weakened. Invasion success drastically increases when we switch the interactions that influence the invader from inhibition to facilitation. Number of instances examined N_s_=1000. Interactions among resident members are equally likely to be facilitative or inhibitory (*f*_*fac*_=0.5). Normalized basal growth rate of the invader is 1.5. Normalized introduced propagule size is 0.3%. The amount of the limiting resource is set at 10^6^ fmole/ml.

**Fig S11.**
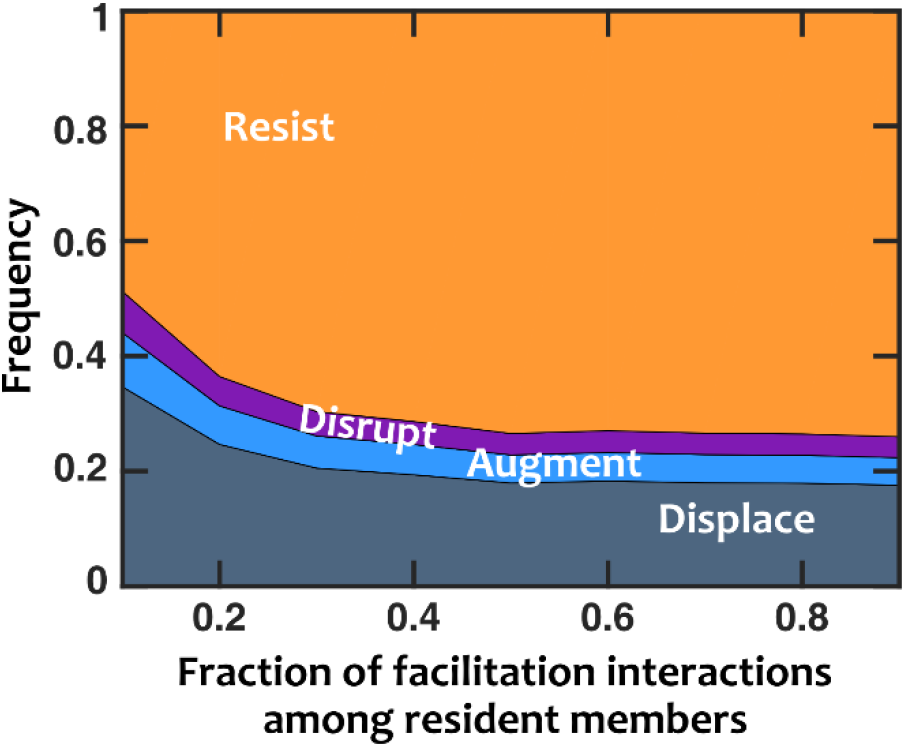
Facilitation among resident species strengthens colonization resistance, even with explicit resource competition. Invasion success decreases when interactions among resident species are predominantly facilitation rather than inhibition. The interactions between resident species and the invader are mostly inhibitory (*f*_*fac,inv*_=0.1). Normalized basal growth rate of the invader is 1.5 (compared to resident members). Normalized introduced propagule size is 0.3%. The amount of the limiting resource is set at 10^6^ fmole/ml.

